# Chronic infections can shape epidemic exposure: Pathogen co-occurrence networks in the Serengeti lions

**DOI:** 10.1101/370841

**Authors:** Nicholas M. Fountain-Jones, Craig Packer, Maude Jacquot, F. Guillaume Blanchet, Karen Terio, Meggan E. Craft

## Abstract

Pathogens are embedded in a complex network of microparasites that can collectively or individually alter disease dynamics and outcomes. Chronic pathogens, for example, can either facilitate or compete with subsequent pathogens thereby exacerbating morbidity and mortality. Pathogen interactions are ubiquitous in nature, but poorly understood, particularly in wild populations. We report here on ten years of serological and molecular data in African lions, leveraging comprehensive demographic and behavioral data to utilize pathogen networks to test if chronic infections shape infection by acute pathogens. We combine network and community ecology approaches to assess broad network structure and characterize associations between pathogens across spatial and temporal scales. We found significant non-random structure in the lion-pathogen co-occurrence network and identified potential facilitative and competitive interactions between acute and chronic pathogens. Our results provide a novel insight for untangling the complex associations underlying pathogen co-occurrence networks.

## Introduction

Identifying and determining the nature of species interactions of multiple pathogens within hosts is increasingly considered critical to understanding infectious disease dynamics (e.g., Pedersen & Fenton 2007; Johnson *et al.* 2015; Seabloom *et al.* 2015; Fountain-Jones *et al.* 2018). Individuals are often co-infected by a diverse infra-community of pathogens, and interactions between pathogens can both alter infection dynamics (Cattadori *et al.* 2008; Susi *et al.* 2015) and influence disease outcomes (Moss *et al.* 2008; Munson *et al.* 2008; Wejse *et al.* 2015). Pathogens that form chronic, long-lasting infections are ubiquitous and are likely to overlap pathogens that produce short-lasting acute infections (Fenton 2008; Randall *et al.* 2013; Rynkiewicz *et al.* 2015; Aivelo & Norberg 2018). Chronic pathogens can compete for the same resources as acute pathogens and can reduce the likelihood of infection (Randall *et al.* 2013; Budischak *et al.* 2018) or, conversely, facilitate infection via immune suppression (e.g., Geldmacher & Koup 2012). Interactions between chronic pathogens, such as between helminths and bacteria, are relatively well understood (e.g., Ezenwa 2016), yet interactions between chronic and acute pathogens, particularly between microparasite species, are largely unknown. As humans, for example, are on average exposed to ten chronic and acute virus species alone (Xu *et al.* 2015), untangling the ecological interactions between microparasites represents a clear knowledge gap that has significant consequences for understanding pathogen dynamics (Munson *et al.* 2008).

Discriminating between positive or negative interactions between pathogens in populations is complicated by the short time window that an acute pathogen is shedding (and thus detectable with molecular methods) and by potentially confounding host environments (Tompkins *et al.* 2011). This is particularly the case for microparasites where pathogen detection often relies on serology, and, thus, without resampling the same individual, the precise timing of exposure cannot be estimated. Detection of chronic pathogens may be more straightforward as the infection is active for more extended periods, but deducing pathogen interactions is difficult without extensive longitudinal data (Tompkins *et al.* 2011; Fenton *et al.* 2014). Identifying whether two pathogens interact due to host-habitat preferences, the increasing likelihood of exposure with age, or infra-community processes such as competition is methodologically challenging (Poulin 2007; Johnson & Buller 2011; Fenton *et al.* 2014; Clark *et al.* 2016).

Further, competition between pathogens could result in increased mortality associated with a particular combination of pathogens (Su *et al.* 2005).

Detecting interactions between pathogens is also likely to depend on taxonomic and spatial scales that are seldom considered (Stutz *et al.* 2018). Studies commonly aggregate pathogen data to genus level, but interactions between pathogens can be subtype or genotype-specific (e.g., Wejse *et al.* 2015; Benesh & Kalbe 2016; Brook *et al.* 2017). For example, individuals infected with human immunodeficiency virus subtype 1 (HIV-1) are four times more likely to become co-infected with tuberculosis compared to individuals with HIV-2 (Wejse *et al.* 2015). Beyond subtype or genus, genotype-specific associations have been demonstrated in snails infected by trematodes (Louhi et al. 2015) and in rodents infected by *Bartonella* bacteria (Brook et al. 2017). Infra-community dynamics are also likely to vary with spatiotemporal scale. For example, pathogen associations in amphibian communities differed substantially at the host scale compared to the pond scale, but these effects can be reversed or undetected at other scales (Stutz et al. 2018). In general, interactions between free-living species are more apparent at scales where the species actually interact compared to larger spatiotemporal scales (Araújo & Rozenfeld 2014).

Recent advances in network theory and community ecology provide an exciting opportunity not only to detect complex interactions between pathogens but also measure the degree that these interactions result from scale-dependent environmental and host factors such as age (Clark *et al.* 2016; Aivelo & Norberg 2018; Stutz *et al.* 2018). Network measures have frequently been used to study food webs but are increasingly applied to pathogen infra-communities where edges represent pathogen co-occurrences within the host (Griffiths *et al.* 2014; Vaumourin *et al.* 2015). Networks are ‘nested’ if pathogens frequently share interaction partners, or ‘segregated’ or modular if the inverse is true. Nested architecture is thought to promote species coexistence if the network is primarily mutualistic (e.g., plant-pollinator networks), but may cause instability if the network is antagonistic (e.g., plant-herbivory networks; Thebault & Fontaine 2010; Rohr *et al.* 2014; Strona & Veech 2015). If the network includes both mutualistic and antagonistic interactions, the effects on the stability of each type of network architecture may be weakened (Sauve *et al.* 2014). Significant segregation within the network also provides evidence for network stability with more modular networks potentially more stable to perturbation (Thebault & Fontaine 2010; Griffiths *et al.* 2014). Understanding network structure could have implications for pathogen control as, for example, if networks are highly segregated and modular, targeted control of one ‘keystone’ pathogen may lead to co-extinction of other pathogens (Pedersen & Fenton 2007; Säterberg *et al.* 2013). Conversely, a nested but unstable network structure could underpin why, for example, control of helminth populations can increase the susceptibility of a host to another infection (Harris et al. 2009).

Although pathogen co-occurrence networks are valuable for quantifying broad structural patterns, they do not account for environmental or host factors or differences in spatial or temporal scale. Joint species distribution models fill this gap by simultaneously assessing environmental influences and interspecific co-occurrences across multiple scales using hierarchical Bayesian mixed models (Warton *et al.* 2015; Ovaskainen *et al.* 2017). Here we use both co-occurrence network and joint species distribution models to examine the structure of pathogen-pathogen networks and quantify pathogen associations whilst controlling for environmental and host factors. We collate ten years of data on acute and chronic pathogens in 105 African lions (*Panthera leo*) as well as extensive host and environmental data from the Serengeti Lion Project (SLP, Packer *et al.* 2005). The SLP datasets provides a unique opportunity to understand pathogen co-occurrence networks in a wild population whilst controlling for group, individual and environmental characteristics. We use this data to ask the following interlinked questions:

(I) To what degree is the co-occurrence network of pathogens in Serengeti lions nested or segregated?
(II) After accounting for environmental and host factors, is the type of chronic infection an individual is infected by associated with subsequent exposure to an acute pathogen? Are there important chronic-chronic or acute-acute pathogen co-occurrence patterns?

We view co-occurrence at two levels of taxonomic resolution and across spatiotemporal scales. We describe a step-analytical pathway that can assess broad network structure and quantify pathogen interactions across multiple scales that can generally be applied to infectious disease dynamics. Detecting pathogen co-occurrence patterns not only provides novel insights into pathogen infra-community dynamics but also helps aide surveillance efforts and generates testable hypotheses that can be answered in laboratory experiments.

## Methods

### Pathogen data

Serological testing and quantitative PCR (qPCR) was performed to detect acute and chronic pathogens from blood samples taken from lions in the Serengeti National Park, Tanzania from 1984-1994. In total, 394 individuals were sampled throughout this period, but our analysis was restricted to the 105 individuals tested for the full suite of pathogens (see Table 1 for pathogen natural history and Table S1 for the number of individuals tested per year included in the analyses). Nomadic individuals (i.e. lions that were not resident in any pride) were excluded due to the difficulty of assigning environmental variables (see *Confounding variables* below). Serological data for canine distemper virus (CDV), feline calicivirus, parvovirus, and coronavirus has been published previously (Packer *et al.* 1999 but see Table S2 for assay details), with the exception of Rift Valley Fever (RVF). To detect RVF exposure we conducted a plaque reduction virus neutralizing test (PRNT) that quantified virus neutralizing antibodies from serum as per the Scott *et al.* (1986) protocol. Anthrax (*Bacilus anthracis*) and feline herpes virus were excluded from analysis because all lions were seropositive (Packer *et al.* 1999; Lembo *et al.* 2011). Similarly, feline leukemia virus (FeLV) was also excluded because al lions were seronegative (Hofmann-Lehmann *et al.* 1996).

**Table 1:**
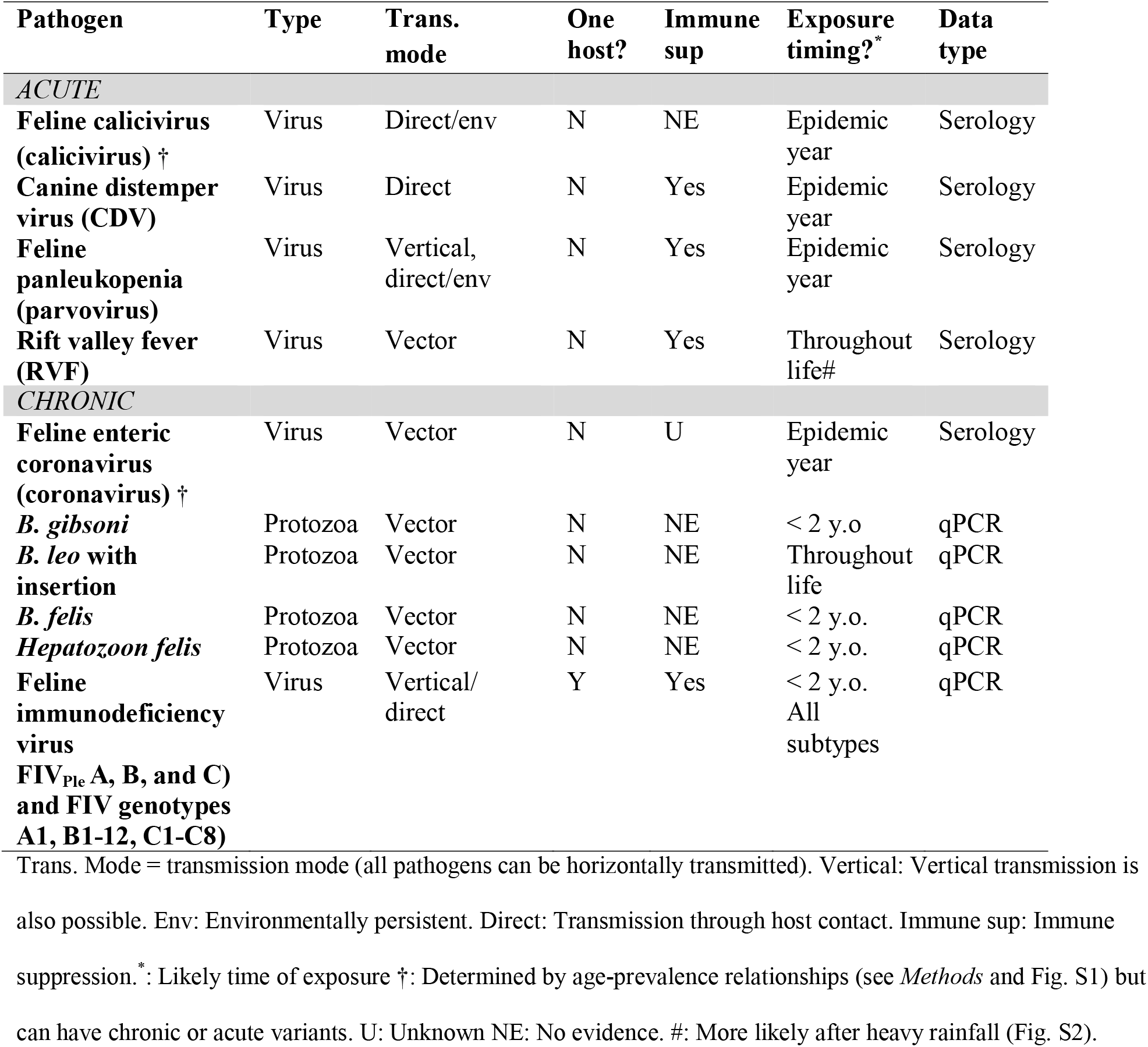
Natural history of both acute and chronic pathogens in this study.

We used qPCR for identification of nucleotides for feline immunodeficiency virus (FIV_Ple_) and the protozoan pathogens in this study (Table 1). Three distinct subtypes of FIV_Ple_ co-circulate in lions in the Serengeti (Troyer *et al.* 2005, 2011; Antunes *et al.* 2008) and thus subtype specific qPCR was performed (see Troyer *et al.* 2004, 2005 for qPCR protocols). The resultant 300 base pair sequences from the *pol* gene were aligned and assigned to 21 operational taxonomic units/genotypes based on a 95% molecular similarity threshold (see Fountain-Jones *et al.* 2017 for details). Lions also commonly get chronically infected by a rich protozoan fauna including *Babesia* and *Hepatazoon* genera. We developed quantitative PCR protocols using density gel gradient electrophoresis to identify each protozoan species (see Munson *et al.* 2008).

We categorized each pathogen as chronic or acute based on the literature and, as the biology of most of these pathogens in lions are poorly understood, age-prevalence relationships. Pathogens which peak in prevalence at a young age (<= 2 y.o.) with little temporal fluctuation were considered to be chronic, whereas an increasing age-prevalence relationship and high temporal variation were classified as more acute (Fig. S1). Feline coronavirus can have both acute and chronic forms and it is impossible to assess which form the individual was exposed to from serological data, but based on age-prevalence relationships we categorized it as a chronic infection (Fig. S1). As most individual lions were likely to be infected by chronic infections within the first two years after birth (Troyer et al. 2011, Fig. S1), we assume that acute exposure typically occurred whilst the individual was infected by the chronic pathogen. We then partitioned the chronic pathogen data into two sets based on taxonomic resolution (high and medium). The high taxonomic resolution dataset encompassed FIV_Ple_ genotype and *Babesia* species data, whereas the medium resolution dataset aggregated FIV_Ple_ subtype information and *Babesia* data to genus level.

### Summary co-occurrence network

We created a summary co-occurrence matrix (n*n) (Griffiths *et al.* 2014) that described the amount of co-occurrence of both chronic and acute pathogens within individuals by multiplying the incidence matrix (i.e., contingency table that described the occurrence of pathogens across individuals) by its transpose. We then computed a measure (□ □) of network structure and modularity index based on node overlap and segregation (Strona & Veech 2015). □ □ is scaled from −1 to 1, where −1 represents an entirely segregated network and 1 represents an entirely nested network.

The co-occurrence matrix was also used to evaluate preferential associations among pathogens utilizing a modularity-based ‘‘greedy’’ approach (Clauset *et al.* 2004). Measures of modularity aim to determine the adequacy of different classification schemes in representing clusters and divisions in datasets; here, the clusters represented the co-occurrence of pathogens in individual hosts. Estimates of modularity were calculated for each possible classification by comparing the expected fraction of co-occurrence of pathogens compared to the quantity if the co-occurrences happened randomly (Newman 2006). The classification with the highest modularity from all the generated classifications was selected. This analysis was performed using the igraph library in R (Csárdi & Nepusz 2006).

The co-occurrence matrix was obtained from the incidence matrix using the graph.incidence and the bipartite.projection functions. The classification analysis was performed using the fastgreedy.community function and graphical output were produced using the tkplot function in igraph.

### Joint species distribution modeling

We fitted joint species distribution models (JDSM) for both high and medium taxonomic resolution datasets, combining information on environmental and host covariates (see Confounding variables below for details), to the occurrence data for each of the pathogens. Pathogens detected fewer than five times were excluded from this analysis. We fitted all the JSDMs with Bayesian inference, using “Hierarchical Modelling of Species Communities” (“HMSC” Blanchet *et al.* 2018). For each analysis, we modeled the response pathogen co-exposure matrix using a probit model. In each model we added individual, pride-year (i.e., which pride and year the individual was sampled in) and year sampled as random effects. We utilized the default priors of the framework (described in full detail in Ovaskainen *et al.* 2017) and ran the HMSC model twice using 3 million MCMC samples (the first 300 000 of which being burn-in). Each run was carried out using a different seed. Visual inspection of MCMC traces were performed to assess convergence. In addition, we made sure that the effective sample size (ESS) of each parameter was > 200.

### Confounding variables

As part of the SLP, most of the individuals in this study have been regularly observed since birth (Mosser & Packer 2009). We selected 13 predictor variables that we thought were likely to be important for pathogen exposure and thus could confound possible interaction patterns (Table 2). We included variables that captured individual variability (e.g., age at sampling), and pride characteristics including environmental variables (e.g., average vegetation cover of the pride’s territory; see Table 2 for measurement details). We attempted to calibrate each predictor set based on possible epidemic year (e.g., calculating rainfall for the year the individual was co-infected by the acute pathogen) as determined by Packer *et al.* (1999), however this removed a large number of males from the dataset as their natal pride were unknown and this reduced the power of our analyses.

**Table 2:** Details of the individual, pride-level and environmental predictors used in the joint species distribution models to help account for potential confounding factors. All variables were calculated based on the year of sampling.

| Predictor | Type | Measurement details | Data |
| --- | --- | --- | --- |
| Sex | Individual | Male or female. | SLP data |
| Age | individual | Age (in days) when lion was sampled | SLP data |
| Number of immigration | Individual | Number of prides an individual has immigrated into prior to sampling. | SLP data |
| Pride, coalition male or nomad? | Individual | Was the individual pride-living or nomadic in the previous two years* prior to exposure of infection (binary). | SLP data |
| Group size | Pride | Average number of individuals in pride two years<sup>†</sup> prior sample collection. | SLP data |
| Despotic | Pride | Was the pride considered despotic at time of sample collection? | SLP data |
| Territory size | Pride | Based on location data over a two-year period based on utilization–distribution curves with a 75% kernel. | SLP data |
| Territory overlap | Pride | What percentage of pride territory overlapped with other prides. | SLP data |
| Habitat quality | Pride | Pride habitat quality score calculated across a two-year period. | (Mosser *et al.* 2009) |
| Number of neighbors | Pride | Number of individuals in neighboring prides. Neighboring prides had territory overlap. | SLP data |
| Yearly rainfall | Environmental | Yearly rainfall experienced in each pride territory based on weather stations in the plains and woodlands. | (Sinclair *et al.* 2013) |
| Average vegetation cover | Environmental | Average vegetation cover across the prides territory based on a 75% kernel. | (Reed *et al.* 2009) |
| Soil Ph | Environmental | Average pH throughout the prides territory base on a 75% kernel. | World Harmonized Soil Database (FAO & IIASA 2009). |

\* We calculated this predictor two years prior to sampling to account for differences in individual status at a potential time of exposure or infection (e.g., individuals that had just immigrated into a pride when sampled were considered nomads as exposure or infection was likely to have occurred previously. †: We averaged over two years to reduce the variability in pride counts as exposure was unlikely to have happened during the sampling year.

## Results

Not including anthrax and herpes (to which all individuals had been exposed), the Serengeti lions were exposed to an average of 5 pathogens (two acute and three chronic, SD = 1) with one individual had been infected by 9 of 10 pathogens (based on medium resolution data, Fig. S3). Remarkably, cubs between 1 and 2 y.o. had been already infected with an average of 4 pathogens (SD = 1) with one 1.5 y.o cub positive for 5. All lions were qPCR positive for at least one protozoan species, and 25% of them were infected by all four considered.

### Pathogen co-occurrence networks are highly nested and modular

The high taxonomic resolution summary network indicated a significantly nested architecture (□ □ = 0.74) with relatively low modularity (modularity index = 0.393, z = 3.307, p = <0.001) with three clusters (Fig. 1a). The largest cluster (green nodes) included all of the protozoans, acute pathogens, and some FIV_Ple_ genotypes, whilst the remaining two clusters consisted just of FIV_Ple_ genotypes (Fig. 1a). While the high prevalence FIV_Ple_ genotypes clustered in the largest cluster, the B2, B3 and C2 genotypes clustered separately, and the disconnected C7 was only detected in one pride during the 1980s. The light red cluster is likely to result from the restricted distribution and temporal period in which these genotypes were identified (e.g., the north western region of the SLP in the 1990’s). The blue and green clusters, in contrast, included a variety of prides over the full sampling period (with the exception of B2 and B6, which were only detected in the woodlands in the 1980’s). Phylogenetically similar genotypes of FIV did not cluster together (Fig. S4). In contrast, the medium resolution network was completely nested with no modularity (□ □ = 1, modularity index = 0, z = ∞, p = 0) and no significant clusters (Fig. 1b).

**Fig. 1:**
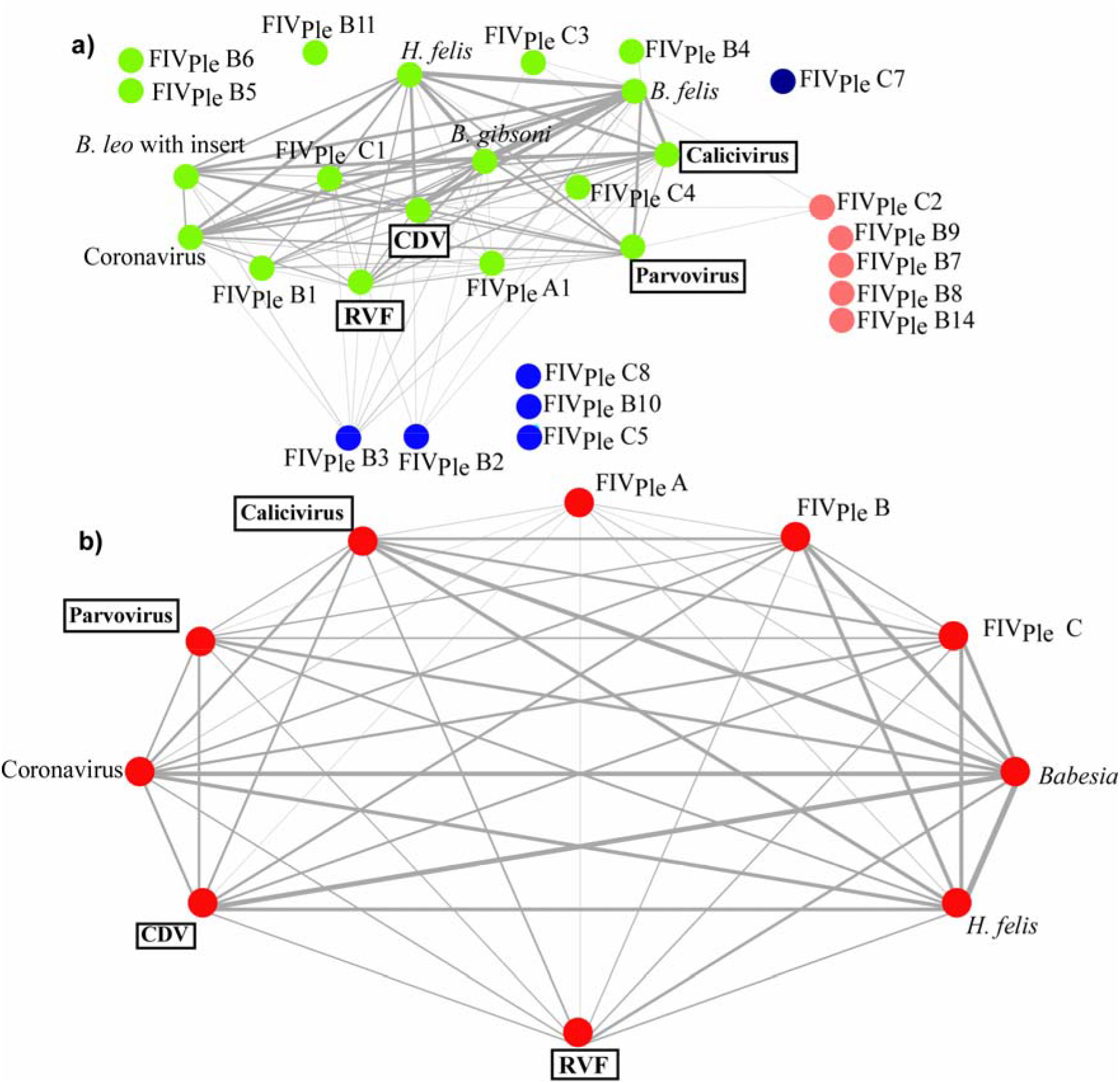
Pathogen summary co-occurrence network for a) high taxonomic resolution and b) medium taxonomic resolution data, where nodes are pathogens and edges reflect co-occurrence. Edges are shown only when there were ≥ 3 co-occurrences. Distinct co-occurrence clusters were identified using a “greedy approach” (Clauset *et al.* 2004); node colors reflect separate clusters. Edge weights are proportional to the number of co-occurrences.

### Strong associations between acute and chronic pathogens across scales

After accounting for environmental, individual and pride factors, the JSDM analysis identified strong associations between pathogens (Fig. 2) that were not detected in the summary co-occurrence network. The nature of these interactions, however, was dependent on the scale assessed. At an individual and pride-year scale, we found strong associations between a small subset of acute and chronic pathogens. FIV_Ple_ B and *H. felis* were negatively associated with RVF (Fig. 2), and FIV_Ple_ B was also negatively associated with parvovirus (Fig. 2a/S5a). However, this interaction could only be detected at medium taxonomic resolution. In contrast, at high taxonomic resolution, we identified positive interactions between *B. gibsoni* and RVF that were not detected at medium-resolution.

**Fig. 2:**
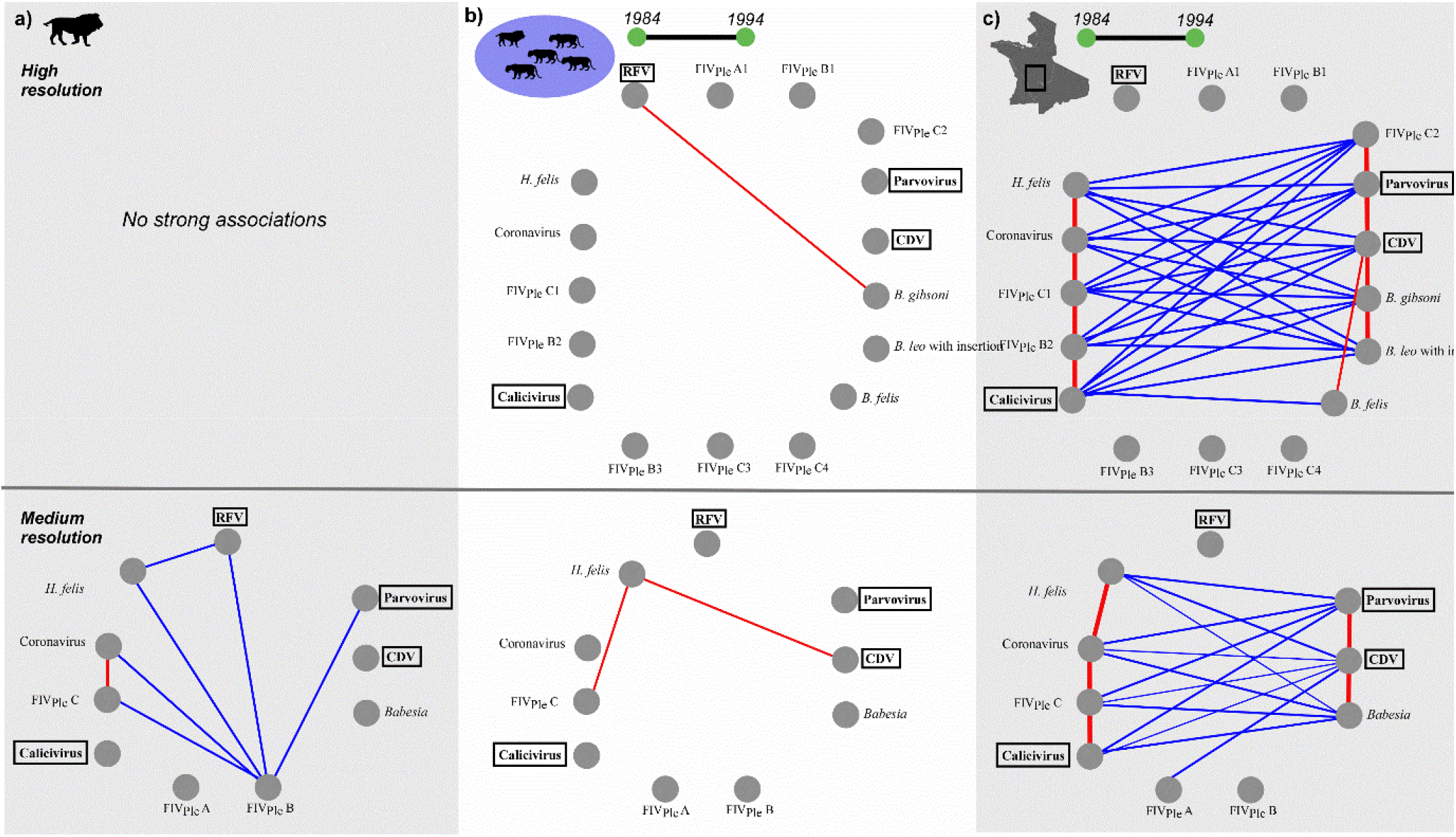
Pathogen-pathogen associations detected at (a) individual, (b) pride-year and (c) landscape-year level after controlling for individual, pride, and environmental variables. Blue represents negative correlations and red indicates positive associations. Only associations with posterior coefficient estimate ≥ 0.4 with 95% credible intervals that do not cross 0 are shown. Pathogens in bold (in boxes) are the acute viruses (all other pathogens are chronic). Length of the dark grey borders reflects the number of strong associations with other pathogens. This figure was drawn using the R package ‘circleplot’ (Westgate 2016). See Fig. S5 for association matrices and Figs. S7/S8 for covariate partitioning and effect size.

The strongest associations between chronic and acute pathogens were detected at the lowest spatial-temporal resolution (landscape-year). In the high taxonomic resolution model, pathogens separated into two groups with each group having a very similar association profile. One group was characterized by positive associations between the *Babesia* species, FIV_Ple_ C2, CDV, and parvovirus. The other group was characterized by positive associations between two FIV_Ple_ genotypes (C1 & B2), coronavirus, *H. felis* and calicivirus (Figs. 2c/S5c). There were strong negative associations between pathogens in each separate group (e.g., CDV and FIV_Ple_ C1). Generally, the same associations held in the medium taxonomic resolution models (Fig. 2c/S5c) but with exceptions. For example, FIV_Ple_ C1 and C2 had opposing association profiles, but as FIV_Ple_ C1 had a higher prevalence (Fig. S6), C1 had the same overall association profile as FIV_Ple_ C.

Associations between acute pathogens were rare. At the year-level, we detected positive associations between CDV and parvovirus with both pathogens negatively associated with calicivirus (Fig. 2c/S5c). In contrast, associations between the chronic pathogens were common, but the nature of the interaction also differed at each taxonomic scale. For example, in the medium resolution model, we detected a positive association between *H. felis* and FIV_Ple_ C that was not found in the high-resolution model indicating that FIV_Ple_ subtype but not genotype was important for this association (Fig. 2b/S5b). Strikingly, we found that FIV_Ple_ subtypes had contrasting association profiles. At the individual level, FIV_Ple_ B and C were negatively associated with each other and negatively and FIV_Ple_ C was positively associated with coronavirus whilst FIV_Ple_ B was negatively associated with coronavirus (Fig. 2a/S5a). See Fig. 3 for a summary of all of the associations detected across scales and Figs. S7/8 for model details.

## Discussion

Here we demonstrate non-random pathogen interactions infecting wild African lions, with both facilitative and competitive associations detected between chronic and acute pathogens across different spatiotemporal scales. We found that whilst there was minimal structure in the summary co-occurrence network (Fig. 1a), we uncovered structure after accounting for scale and controlling for potentially confounding environmental and host variables (Fig. 2). The particular chronic pathogen an individual is infected by as a cub may have consequences for which acute pathogen the individual is infected with later in life (Fig. 3). Moreover, we detected strong relationships between chronic pathogens across scales, and these associations may have consequences for lion health. We emphasize that the approach used here can start to untangle pathogen infra-community relationships and identify potential chronic-acute interactions in wild populations. These can then be compared with knowledge of pathogen pathogenesis and validated *in-vitro* in a laboratory setting. Whilst clinical or laboratory studies of co-infection in lions are rare, the associations we found have clear precedence in similar pathogens co-infecting humans. Our results not only provide new insights on disease dynamics in the Serengeti lions but also provide a valuable framework for exploring pathogen co-occurrence networks and infra-community dynamics across taxonomic and spatial scales.

**Fig. 3:**
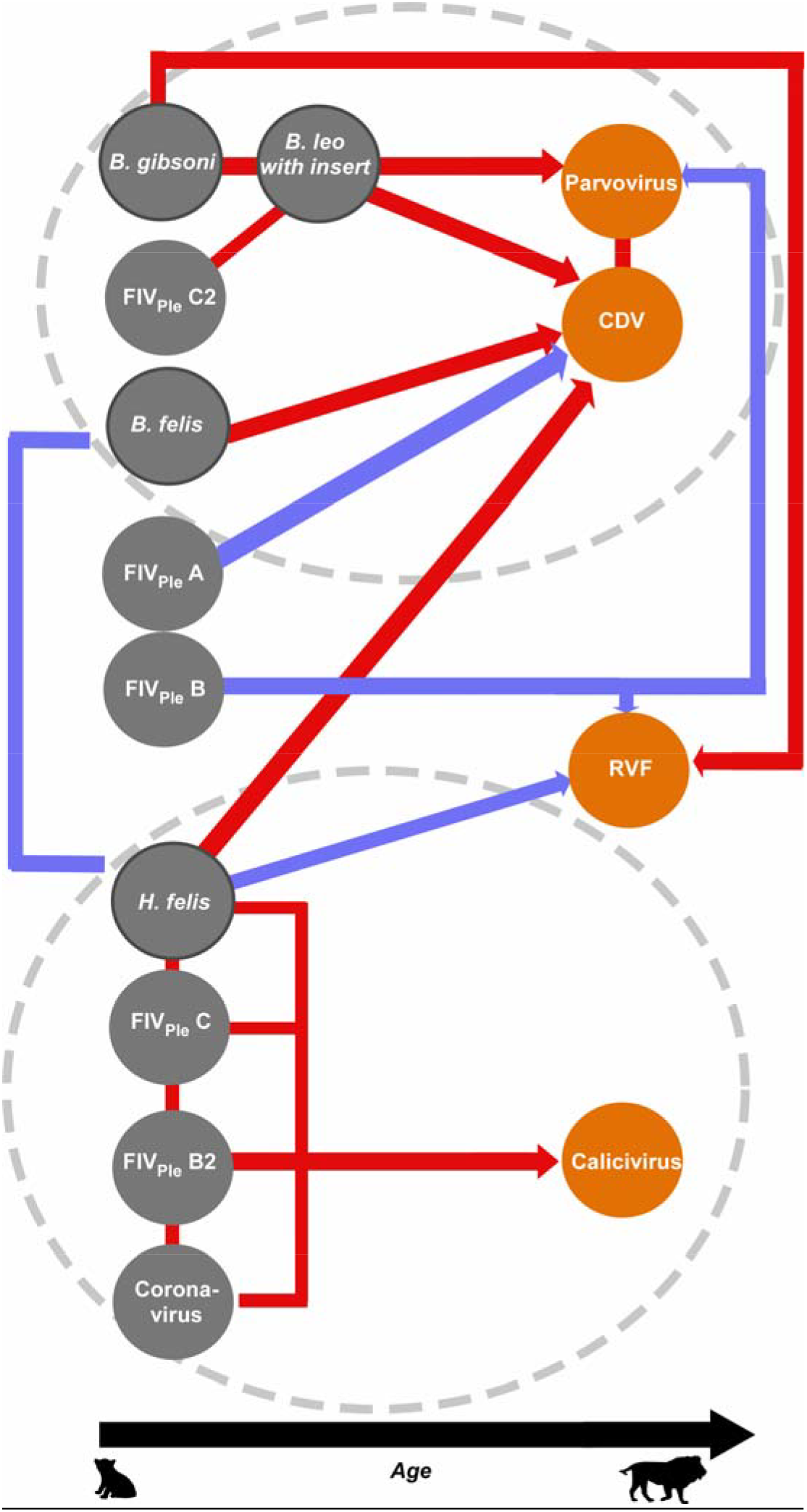
Summary of the strong positive (red line/arrows) and negative (blue lines/arrows) interactions between chronic (grey circles) and acute (orange circles) pathogens in the Serengeti lions; dark-grey borders indicate protozoa. The direction of the red or blue arrows indicates the potential sequence of infection events. The black arrow along the X-axis represents lion age with the relative position of the circles reflecting when lions were likely to be infected by each pathogen (see Fig. S1). Dashed circles indicate major co-occurrence clusters identified at the landscape-year scale.

Co-occurrence networks were highly nested with relatively low modularity, particularly at a medium taxonomic resolution. This contrasts with the more modular human parasite co-occurrence network (Griffiths *et al.* 2014). However, this is not surprising as the human networks included host resources and immune system components as nodes, and we lacked data on bacteria, helminths, and fungi. These additional taxa may lead to further segregation in the network, as larger and more diverse networks typically show increased modularity and segregation (Thebault & Fontaine 2010; Sauve *et al.* 2014). Expanding sampling to construct a more complete microbe and macroparasite network would also capture a broader array of potentially facilitative and competitive interactions (Ezenwa 2016; Aivelo & Norberg 2018).

The predicted stability (or otherwise) of our lion co-occurrence network is uncertain. Theoretically, if such a strongly nested network were mostly mutualistic (i.e., pathogens facilitate other pathogens) if the prevalence of one pathogen decreased significantly due to vaccination, for example, the overall structure would not alter (e.g., Thebault & Fontaine 2010). However, this may not hold for networks consisting of both positive and negative associations as found here (Sauve *et al.* 2014). Conversely, the relatively low segregation in the lion network could mean that a perturbation would be less likely to be restricted to a particular compartment (Griffiths *et al.* 2014). We recommend further empirical and theoretical investigations on the consequences of nested or segregated networks on the stability of pathogen infra-communities.

After accounting for environment and host effects, we found that the chronic pathogens were strongly associated with the acute pathogens across scales. For example, at an individual scale, chronic infections negatively interacted with RVF. Coinfections between bunyaviruses like RVF and retroviruses are likely common in humans and wildlife, though there are surprisingly few studies addressing the topic. In contrast, relationships between dengue virus (a flavivirus) and HIV are relatively well understood. Flaviviruses and HIV share similar immune receptors that can inhibit HIV replication and the molecular machinery used to do so maybe a viable way to control HIV infection (e.g., Xiang *et al.* 2009). Given the overall structural similarity of flaviviruses and bunyaviruses (Hernandez *et al.* 2014), it is possible that a similar mechanism underlies the association in lions between RVF and FIV_Ple_ that we observed, although we show that this association was subtype specific. If this was true, RVF may actually inhibit FIV_Ple_ B infection counter to our assumption that chronic pathogens in our system infected each individual first (Fig. 3).

Relationships between acute and chronic pathogens were strongest at the year-landscape level. The association between CDV and *Babesia* is well documented with high levels of *Babesia* infection magnifying the impacts of consequent T cell depletion caused by CDV infection leading to mortality of nearly 40% of the lion population in 1994 (Munson *et al.* 2008). We found that all tick-borne hemoparasites showed potentially facilitative associations with CDV including *B. leo* (with insert) despite its low prevalence in 1993/4 (Fig. S9) Parvovirus was also positively associated with CDV, but this was likely due to similarities in epidemic cycles with a parvovirus epidemic in 1992 just before the 1994 CDV epidemic (Packer *et al.* 1999) Parvoviruses are also immune suppressive, and so the timing of the parvovirus outbreak may also have contributed to the CDV/*Babesia*-induced mortality. The general negative relationship between FIV_Ple_ C and CDV/*Babesia* supports the theory that individuals infected by subtype C were more likely to die in the consequent *Babesia*/CDV outbreak (Troyer *et al.* 2011). Thus, this negative association may not be due to competition between pathogens but rather to mortality.

Our approach detected strong interactions between the chronic pathogens. For example, there were opposing associations between the FIV_Ple_ subtypes and coronavirus (Fig. 2). Negative associations between retroviruses and coronaviruses are rarely reported, yet there are plausible molecular pathways. HIV-1 and human coronaviruses (HCoV) share remarkably similar binding receptors (Chan *et al.* 2006) and some mild HCoV strains even considered a viable vaccine against HIV (Eriksson *et al.* 2006). This may explain the negative association we detected for FIV_Ple_ B and coronavirus but does not explain the positive association between FIV_Ple_ C and coronavirus we detected across scales. The mechanism driving FIV_Ple_ subtype specific relationships with coronaviruses are unclear, and as coronaviruses infecting lions are also likely to be genetically diverse, examining the genetic structure of coronavirus may help untangle these associations further. In contrast, competitive interactions between HIV strains are well characterized with HIV-1, for example, found to outcompete HIV-2 for blood resources (Ariën *et al.* 2005). For FIV_Ple_, even though co-infection is relatively common (Troyer *et al.* 2011) competition between subtypes could be important as there is anecdotal cell culture evidence that FIV_Ple_ B can propagate more rapidly than FIV_Ple_ C (Melody Roelke, unpublished data).

There were also contrasting associations between the protozoan species. For example, the distribution of *B. felis* was not shaped by any other protozoan and in general, had a narrow association profile (Fig.2), unlike the other *Babesia* species. For the individuals co-infected by protozoans, combinations involving *B. felis* were also common, whereas co-infections involving *H. felis* and the other *Babesia* species varied in prevalence and composition (Fig. S9). Even though *B. gibsoni* and *B. felis* show similar age prevalence profiles (Fig. S1), the prevalence of *B. felis* over time was relatively stable compared to the other protozoa (Fig. S10). Differences in the host range for individual *Babesia* species and potential host differences in virulence may partially explain these patterns. For example, *B. felis* has only ever been detected in felids, whereas *B. gibsoni* has a much broader host range including canids (Penzhorn 2006). More generalist pathogens may have greater pathogenicity as there can be reduced selective restraint on virulence particularly in ‘dead end’ hosts (Woolhouse *et al.* 2001). If more pathogenic species are more likely to form significant associations with other pathogens compared to less virulent pathogens is an open question in disease ecology. Importantly, patterns like these would be missed without incorporating high-resolution pathogen data.

There are, however, limitations to this approach. The inability to distinguish mortality from competitive interactions is one of them, and careful interpretation of negative associations is necessary. Incorporating approaches such as structural equation models that explicitly include potential mechanisms that underly candidate pathogen interactions (Carver *et al.* 2015) could be a valuable additional step in future pathogen network studies. Another weakness is the inability to estimate the timing of these infections more precisely. For example, the negative association between RVF and *H. felis* could be due to differences in rainfall affecting vector abundance with higher rainfall years increase mosquito abundance thus increasing RVF prevalence (Fig. S2). In contrast, low rainfall years increase the risk of lions being exposed to ticks (Munson *et al.* 2008) thus potentially increasing *H. felis* prevalence. As rainfall was calibrated to the age of sampling rather than the age of infection (which could differ) the JSDM approach could not capture this variation. Furthermore, we cannot quantify the importance of these associations in shaping pathogen distribution across scales compared to processes such as host density. Also, the number of samples we had did differ across years (Table S1) with most coming between 1985 and 1994, and this have influenced some patterns, particularly between genotypes. Lastly, incorporating immune function and host resources in both the summary network and JSDM analyses are likely to provide mechanistic insight into pathogen network structure (Griffiths *et al.* 2014). However, given the daunting complexity of pathogen infra-community dynamics, our two-step approach can assess broad network structure and identify useful candidate interactions between pathogens thereby reducing some of this complexity.

The high frequency of co-occurrence and co-infection in lions – and the potential for specific associations to cause population decline – highlights the importance of understanding pathogen associations. The summary lion co-occurrence network is highly connected with both positive and negative associations between chronic and acute pathogens and altering the network could assist with disease control. For example, individuals vaccinated for both parvovirus and *H. felis* could potentially be at lower risk of *Babesia* and CDV infection. Conversely, our networks show that there could be unintended consequences of controlling a pathogen, as has been shown for deworming (Fenton 2013). For example, it is possible that vaccination targeting coronavirus in this population could increase *B. gibsoni* infections that may, in turn, lead to a higher risk of CDV exposure. Whilst these relationships are correlative rather than mechanistic, it is clear from our study that relationships between microparasites are common and should be considered in disease management.

Our findings indicate that the lion’s pathogen infra-community is complex, but interactions between pathogens are important. We identify useful candidate interactions between pathogens thereby reducing some of this complexity. More broadly, our work demonstrates how different network approaches can be combined to gain insights into the ecological factors underlying pathogen structures and interactions and how this can be applied to the study of pathogen communities of wildlife populations. In addition to these biological insights, the study highlights several critical areas for methodological improvement that can currently limit robust inference of pathogen interactions from time-series serological and qPCR data. Addressing these limitations is timely, given the ongoing threat of wildlife populations decline, creating an urgent need to integrate better molecular, ecological and network information to guide strategies for disease control.

## Acknowledgments

N.M F-J. and MEC were funded by National Science Foundation (DEB-1413925 and 1654609), the University of Minnesota’s Office of the Vice President for Research and Academic Health Center Seed Grant, and the Cooperative State Research Service, U.S. Department of Agriculture, under Project No. MIN-62-098. We also thank Dr. Linda Munson who led the CDV-*Babesia* work and Professor CJ Peters for performing the RVF virus neutralization tests.

Author Contributions
NMFJ and MEC designed the study. CP & KT provided data. NMFJ, MJ & GB conducted statistical analyses. NMFJ wrote the manuscript and all authors contributed to revisions.

## References

Aivelo, T. & Norberg, A. (2018). Parasite-microbiota interactions potentially affect intestinal communities in wild mammals. J. Anim. Ecol., 87, 438–447.

Antunes, A., Troyer, J.L., Roelke, M.E., Pecon-Slattery, J., Packer, C., Winterbach, C., et al. (2008). The evolutionary dynamics of the lion *Panthera leo* revealed by host and viral population genomics. PLoS Genet., 4, e1000251.

Ariën, K.K., Abraha, A., Quiñones-Mateu, M.E., Kestens, L., Vanham, G. & Arts, E.J. (2005). The replicative fitness of primary human immunodeficiency virus type 1 (HIV-1) group M, HIV-1 group O, and HIV-2 isolates. J. Virol., 79, 8979–90.

Benesh, D.P. & Kalbe, M. (2016). Experimental parasite community ecology: intraspecific variation in a large tapeworm affects community assembly. J. Anim. Ecol., 85, 1004–1013.

Blanchet, F.G., Tikhonov, G. & Norberg, A. (2018). HMSC: Hierarchical Modelling of Species Community. R package version 2.2-1.

Brook, C.E., Bai, Y., Yu, E.O., Ranaivoson, H.C., Shin, H., Dobson, A.P., et al. (2017). Elucidating transmission dynamics and host-parasite-vector relationships for rodent-borne *Bartonella* spp. in Madagascar. Epidemics, 20, 56–66.

Budischak, S.A., Wiria, A.E., Hamid, F., Wammes, L.J., Kaisar, M.M.M., van Lieshout, L., et al. (2018). Competing for blood: the ecology of parasite resource competition in human malaria-helminth co-infections. Ecol. Lett., 21, 536–545.

Carver, S., Beatty, J.A., Troyer, R.M., Harris, R.L., Stutzman-Rodriguez, K., Barrs, V.R., et al. (2015). Closing the gap on causal processes of infection risk from cross-sectional data: structural equation models to understand infection and co-infection. Parasit. Vectors, 8, 658.

Cattadori, I.M., Boag, B. & Hudson, P.J. (2008). Parasite co-infection and interaction as drivers of host heterogeneity. Int. J. Parasitol., 38, 371–380.

Chan, V.S.F., Chan, K.Y.K., Chen, Y., Poon, L.L.M., Cheung, A.N.Y., Zheng, B., et al. (2006). Homozygous L-SIGN (CLEC4M) plays a protective role in SARS coronavirus infection. Nat. Genet., 38, 38–46.

Clark, N.J., Wells, K., Dimitrov, D. & Clegg, S.M. (2016). Co-infections and environmental conditions drive the distributions of blood parasites in wild birds. J. Anim. Ecol., 85, 1461–1470.

Clauset, A., Newman, M.E.J. & Moore, C. (2004). Finding community structure in very large networks. Phys. Rev. E, 70.

Csárdi, G. & Nepusz, T. (2006). The igraph software package for complex network research. InterJournal Complex Syst., 1695.

Eriksson, K.K., Makia, D., Maier, R., Ludewig, B. & Thiel, V. (2006). Towards a coronavirus-based HIV multigene vaccine. Clin. Dev. Immunol., 13, 353–60.

Ezenwa, V.O. (2016). Helminth-microparasite co-infection in wildlife: lessons from ruminants, rodents and rabbits. Parasite Immunol., 38, 527–534.

FAO & IIASA. (2009). Harmonized world soil database. Food Agric. Organ.

Fenton, A. (2008). Worms and germs: the population dynamic consequences of microparasite-macroparasite co-infection. Parasitology, 135, 1545–1560.

Fenton, A. (2013). Dances with worms: the ecological and evolutionary impacts of deworming on coinfecting pathogens. Parasitology, 140, 1119–32.

Fenton, A., Knowles, S.C.L., Petchey, O.L. & Pedersen, A.B. (2014). The reliability of observational approaches for detecting interspecific parasite interactions: comparison with experimental results. Int. J. Parasitol., 44, 437–445.

Fountain-Jones, N.M., Packer, C., Troyer, J.L., VanderWaal, K., Robinson, S., Jacquot, M., et al. (2017). Linking social and spatial networks to viral community phylogenetics reveals subtype-specific transmission dynamics in African lions. J. Anim. Ecol., 86, 1469–1482.

Fountain-Jones, N.M., Pearse, W.D., Escobar, L.E., Alba-Casals, A., Carver, S., Davies, T.J., et al. (2018). Towards an eco-phylogenetic framework for infectious disease ecology. Biol. Rev., 93, 950–970.

Geldmacher, C. & Koup, R.A. (2012). Pathogen-specific T cell depletion and reactivation of opportunistic pathogens in HIV infection. Trends Immunol., 33, 207–14.

Griffiths, E.C., Pedersen, A.B., Fenton, A. & Petchey, O.L. (2014). Analysis of a summary network of co-infection in humans reveals that parasites interact most via shared resources. Proceedings. Biol. Sci., 281, 20132286.

Hernandez, R., Brown, D.T. & Paredes, A. (2014). Structural differences observed in arboviruses of the alphavirus and flavivirus genera. Adv. Virol., 2014, 259382.

Hofmann-Lehmann, R., Fehr, D., Grob, M., Elgizoli, M., Packer, C., Martenson, J.S., et al. (1996). Prevalence of antibodies to feline parvovirus, calicivirus, herpesvirus, coronavirus, and immunodeficiency virus and of feline leukemia virus antigen and the interrelationship of these viral infections in free-ranging lions in East Africa, 3, 554–562.

Johnson, P.T.J. & Buller, I.D. (2011). Parasite competition hidden by correlated coinfection: using surveys and experiments to understand parasite interactions, 92, 535–541.

Johnson, P.T.J., de Roode, J.C. & Fenton, A. (2015). Why infectious disease research needs community ecology. Science (80-.)., 349, 1259504–1259504.

Lembo, T., Hampson, K., Auty, H., Beesley, C.A., Bessell, P., Packer, C., et al. (2011). Serologic surveillance of anthrax in the Serengeti ecosystem, Tanzania, 1996–2009. Emerg. Infect. Dis., 17, 387–394.

Moss, W.J., Fisher, C., Scott, S., Monze, M., Ryon, J.J., Quinn, T.C., et al. (2008). HIV Type 1 infection is a risk factor for mortality in hospitalized Zambian children with measles. Clin. Infect. Dis., 46, 523–527.

Mosser, A., Fryxell, J.M., Eberly, L. & Packer, C. (2009). Serengeti real estate: density vs. fitness-based indicators of lion habitat quality. Ecol. Lett., 12, 1050–1060.

Mosser, A. & Packer, C. (2009). Group territoriality and the benefits of sociality in the African lion, Panthera leo. Anim. Behav., 78, 359–370.

Munson, L., Terio, K.A., Kock, R., Mlengeya, T., Roelke, M.E., Dubovi, E., et al. (2008). Climate extremes promote fatal co-infections during canine distemper epidemics in African lions. PLoS One, 3, e2545.

Newman, M.E.J. (2006). Community structure in social and biological networks. Proc. Natl. Acad. Sci. U. S. A., 99, 7821–6.

Ovaskainen, O., Tikhonov, G., Norberg, A., Guillaume Blanchet, F., Duan, L., Dunson, D., et al. (2017). How to make more out of community data? A conceptual framework and its implementation as models and software. Ecol. Lett., 20, 561–576.

Packer, C., Altizer, S., Appel, M., Brown, E., Martenson, J., O’Brien, S.J., et al. (1999). Viruses of the Serengeti: patterns of infection and mortality in African lions. J. Anim. Ecol., 68, 1161–1178.

Packer, C., Hilborn, R., Mosser, A., Kissui, B., Borner, M., Hopcraft, G., et al. (2005). Ecological change, group territoriality, and population dynamics in Serengeti lions. Science (80-.)., 307, 390–3.

Pedersen, A.B. & Fenton, A. (2007). Emphasizing the ecology in parasite community ecology. Trends Ecol. Evol. Evol., 22, 133–9.

Penzhorn, B.L. (2006). Babesiosis of wild carnivores and ungulates. Vet. Parasitol., 138, 11–21.

Poulin, R. (2007). Are there general laws in parasite ecology? Parasitology, 134, 763–766.

Randall, J., Cable, J., Guschina, I.A., Harwood, J.L. & Lello, J. (2013). Endemic infection reduces transmission potential of an epidemic parasite during co-infection. Proceedings. Biol. Sci., 280, 20131500.

Reed, D.N., Anderson, T.M., Dempewolf, J., Metzger, K. & Serneels, S. (2009). The spatial distribution of vegetation types in the Serengeti ecosystem: the influence of rainfall and topographic relief on vegetation patch characteristics. J. Biogeogr., 36, 770–782.

Rohr, R.P., Saavedra, S. & Bascompte, J. (2014). Ecological networks. On the structural stability of mutualistic systems. Science, 345, 1253497.

Rynkiewicz, E.C., Pedersen, A.B. & Fenton, A. (2015). An ecosystem approach to understanding and managing within-host parasite community dynamics. Trends Parasitol., 31, 212–221.

Säterberg, T., Sellman, S. & Ebenman, B. (2013). High frequency of functional extinctions in ecological networks. Nature, 499, 468–470.

Sauve, A.M.C., Fontaine, C. & Thébault, E. (2014). Structure-stability relationships in networks combining mutualistic and antagonistic interactions. Oikos, 123, 378–384.

Scott, R.M., Feinsod, F.M., Allam, I.H., Ksiazek, T.G., Peters, C.J., Botros, B.A.M., et al. (1986). Serological tests for detecting rift valley fever viral antibodies in sheep from the Nile Delta. J. Clin. Microbiol., 24, 612–614.

Seabloom, E.W., Borer, E.T., Gross, K., Kendig, A.E., Lacroix, C., Mitchell, C.E., et al. (2015). The community ecology of pathogens: coinfection, coexistence and community composition. Ecol. Lett., 18, 401–415.

Sinclair, A.R.E., Metzger, K.L., Fryxell, J.M., Packer, C., Byrom, A.E., Craft, M.E., et al. (2013). Asynchronous food-web pathways could buffer the response of Serengeti predators to El Niño Southern Oscillation. Ecology, 94, 1123–1130.

Strona, G. & Veech, J.A. (2015). A new measure of ecological network structure based on node overlap and segregation. Methods Ecol. Evol., 6, 907–915.

Stutz, W.E., Blaustein, A.R., Briggs, C.J., Hoverman, J.T., Rohr, J.R. & Johnson, P.T.J. (2018). Using multi-response models to investigate pathogen coinfections across scales: Insights from emerging diseases of amphibians. Methods Ecol. Evol., 9, 1109–1120.

Su, Z., Segura, M., Morgan, K., Loredo-Osti, J.C. & Stevenson, M.M. (2005). Impairment of protective immunity to blood-stage malaria by concurrent nematode infection. Infect. Imunity, 73, 3531–9.

Susi, H., Barrès, B., Vale, P.F., Laine, A.-L., Mideo, N., Alizon, S., et al. (2015). Co-infection alters population dynamics of infectious disease. Nat. Commun., 6, 5975.

Thebault, E. & Fontaine, C. (2010). Stability of ecological communities and the architecture of mutualistic and trophic networks. Science (80-.)., 329, 853–856.

Tompkins, D.M., Dunn, A.M., Smith, M.J. & Telfer, S. (2011). Wildlife diseases: from individuals to ecosystems. J. Anim. Ecol., 80, 19–38.

Troyer, J.L., Pecon-Slattery, J., Roelke, M.E., Black, L., Packer, C. & O’Brien, S.J. (2004). Patterns of feline immunodeficiency virus multiple infection and genome divergence in a free-ranging population of African lions. J. Virol., 78, 3777–3791.

Troyer, J.L., Pecon-Slattery, J., Roelke, M.E., Johnson, W., VandeWoude, S., Vazquez-Salat, N., et al. (2005). Seroprevalence and genomic divergence of circulating strains of feline immunodeficiency virus among Felidae and Hyaenidae species. J. Virol., 79, 8282–8294.

Troyer, J.L., Roelke, M.E., Jespersen, J.M., Baggett, N., Buckley-Beason, V., MacNulty, D., et al. (2011). FIV diversity: FIV _Ple_ subtype composition may influence disease outcome in African lions. Vet. Immunol. Immunopathol., 143, 338–346.

Vaumourin, E., Vourc’h, G., Gasqui, P., Vayssier-Taussat, M., Windsor, D., Anderson, R., et al. (2015). The importance of multiparasitism: examining the consequences of co-infections for human and animal health. Parasit. Vectors, 8, 545.

Warton, D.I., Blanchet, F.G., O’Hara, R.B., Ovaskainen, O., Taskinen, S., Walker, S.C., et al. (2015). So many variables: Joint modeling in community ecology. Trends Ecol. Evol., 30, 766–79.

Wejse, C., Patsche, C.B., Kühle, A., Bamba, F.J.V., Mendes, M.S., Lemvik, G., et al. (2015). Impact of HIV-1, HIV-2, and HIV-1+2 dual infection on the outcome of tuberculosis. Int. J. Infect. Dis., 32, 128–134.

Westgate, M.J. (2016). circleplot: Circular plots of distance and association matrices. R package version 0.4.

Woolhouse, M.E.J., Taylor, L.H. & Haydon, D.T. (2001). Population biology of multihost pathogens. Science (80-.)., 292, 1109–1112.

Xiang, J., McLinden, J.H., Rydze, R.A., Chang, Q., Kaufman, T.M., Klinzman, D., et al. (2009). Viruses within the Flaviviridae decrease CD4 expression and inhibit HIV replication in human CD4+ cells. J. Immunol., 183, 7860–9.

Xu, G.J., Kula, T., Xu, Q., Li, M.Z., Vernon, S.D., Ndung’u, T., et al. (2015). Comprehensive serological profiling of human populations using a synthetic human virome. Science (80-.)., 348, aaa0698–aaa0698.

